# Valuing conservation and natural wealth: The blue economy of manta ray watching in the Maldives

**DOI:** 10.1101/2025.06.05.658185

**Authors:** Hannah M. Moloney, Maria I. Garcia Rojas, Nina Rothe, Asia O. Armstrong, Kirsty Ballard, Florence Barraud, Farah Hamdan, Anthony J. Richardson, Enas Mohamed Riyad, Tamaryn J. Sawers, Kathy A. Townsend, Guy M.W. Stevens

**Author notes:** These authors contributed equally to this work (Maria I. Garcia Rojas, Nina Rothe).

## Abstract

Amid declining manta ray populations globally, the well-established and growing manta ray tourism industries generate substantial economic benefits and aid protective legislation for these threatened elasmobranchs. As flagship species, manta rays are a drawcard for marine wildlife tourism and a gateway for engaging the public and communities in conservation. Healthy marine ecosystems are the key drivers of employment and economic sustainability for island nations such as the Maldives. However, there are many stakeholders competing for these shared resources, which can result in environmental degradation. Economic evaluations are a powerful tool for justifying the conservation efforts of threatened species and natural areas, especially in light of competing stakeholders. Using tour operator surveys (*n*=106) and data mining, this study provides an updated assessment of manta ray watching tourism in the Maldives, and represents the first national evaluation of its total economic and socio-economic benefits. In 2021, manta ray tourism in the Maldives generated an estimated US$227.3 million, including US$39 million on manta ray focused diving and snorkelling excursions, and US$188.3 million in related tourist expenditure. This represented 4.8% of the national Gross Domestic Product. This industry appears to have grown around 380% since 2008 (US$8.1 million) and manta ray watching is now offered by 80% of tourism operators nation-wide. Our findings revealed that manta rays hold intrinsic value and cultural significance within local communities. Acknowledging this, the flow-on benefits to the community extend beyond this industry, reaching local businesses, employed staff, and the government with the total economic benefits of the manta ray tourism industry were estimated at over US$311 million per year. Such value highlights the significance of manta rays for this nation and the need for effective management centred on manta ray conservation to safeguard future prosperity and mitigate the potential impact of tourism on manta ray populations.

## Introduction

Wildlife tourism can support both economic growth and the conservation of natural resources and ecosystems [1–4]. Wildlife viewing activities are a cultural ecosystem service due to the non-material benefits obtained from ecosystems through recreation, aesthetic spiritual experiences, learning and cultural enrichment [5]. These ecosystem services are based on both human residents and visitors valuing the intrinsic beauty of wild species and places, providing a motivation for tourists to visit these ecosystems, and for the local community to protect them [5]. Despite this, policy makers are challenged with balancing the needs of multiple stakeholders competing for the use of environmental resources [6]. Highlighting both the short- and long-term social and economic benefits of a profitable wildlife tourism industry can help strengthen the case for conserving threatened species and ecologically important habitats by showcasing the social and economic value of wildlife tourism [7–9]. In well-managed situations, visiting tourists benefit from an enhanced sense of personal wellbeing, as well as increased educational and recreational values [5,10]. For local communities, the economy can be boosted through employment and income generation [9]. These wildlife-tourist interactions can also encourage greater conservation efforts by both operators and visitors [11–14]. Thus, wildlife tourism offers an effective incentive for governments to establish protection measures and encourage stakeholder support [15,16].

Implementing ongoing management and monitoring of protected natural resources and biodiversity can be costly for governments (i.e., staff salaries, governance structure and facilities) [6,17]. To maximize conservation outcomes within a limited budget, the use of flagship species has proven to be an effective approach for the restoration and conservation of biodiversity across the globe [18,19]. The selection of a species for flagship status must be well balanced and consider not only the intrinsic and ecological aspects (e.g., risk of extinction), but also the cultural, social and economic context of the region where the flagship species will be promoted [20,21]. As such, tourism in the marine environment focusing on a flagship species can make important contributions to biodiversity conservation and protect lesser-known species [19]. Marine megafauna tourism has been growing in popularity since the 1980s [1,15,22,23], as have the associated economic benefits, both directly for operators and indirectly to the governments and businesses within local economies [4,24–26]. Notably, the development of well-managed megafauna tourism has provided the opportunity for communities, particularly in remote locations where there are few alternatives, to use their natural resources in a sustainable manner [23,27].

Shark and ray watching is a rapidly growing area of marine wildlife tourism, with opportunities available in at least 42 countries, primarily distributed in low- to middle-income nations, including island nations [3,8,14,28]. Typically, the tourism industry can generate far greater economic returns than fisheries on the focal species [3,13,28]. Indeed, Anderson & Ahmed [29] in 1993 calculated that the tourism value of a live shark over its lifetime in the Republic of Maldives, was US$33,500, compared to US$32 for a dead shark that was fished. Coral-reef associated shark and ray species are among the most threatened marine groups, with two-thirds (59%) of species threatened with extinction according to the International Union for Conservation of Nature (IUCN) Red List of Threatened Species [30]. Many shark and ray populations are experiencing substantial population declines largely due to overfishing, intensified by climate change and habitat degradation [30,31]. Shark and ray tourism can provide an alternative to extractive practices in certain regions of the world where tourism infrastructure and markets exist. Implemented in 29 countries, shark and ray sanctuaries – typically prohibiting all commercial shark and/or ray fishing, trade, possession or sale within their Exclusive Economic Zone – provide an example of a limit-based measure for conservation [32–34]. These sanctuaries have typically been declared in small-island nations, particularly in the tropical Pacific Ocean, the Caribbean Sea and the Indian Ocean [34], where there is a high reliance on marine resources [35] and often a limited capacity for enforcement [36]. Viewing marine megafauna such as sharks and rays as economically important industries led the Maldives’ government to implement a shark sanctuary in 2010 [37,38]. Although rays and skates were first protected from the export trade by the Ministry of Fisheries in 1995/6, it was not until 2014 that the EPA legislation ((IUL)438-ECAS/438/2014/81) made it illegal to kill, catch or harm all rays and skates; effectively making the Maldives both a shark and ray sanctuary [38].

Manta rays are zooplanktivorous giants found throughout tropical and subtropical oceans [39]. They aggregate seasonally at predictable locations that facilitate socialising and key life history functions including feeding, courtship and mating, predator avoidance, cleaning and thermoregulation [40]. The predictable and reliable nature of these aggregations makes manta ray watching (MRW) a highly valued tourist activity [41–43] involving recreational diving, snorkelling, and on-water observation of these animals in the wild [3]. This high value is evident in the Maldives, where tourists are willing to pay more for manta ray encounters than for those with sea turtles and reef sharks [44]. In 2013, MRW was a rapidly growing industry in >25 countries worldwide, valued at US$140 million per year (i.e., tour operator revenue and tourist expenses), with hotspots in the Maldives, Japan, Mozambique, Australia, Hawaii and Indonesia [3]. The Maldives had the largest MRW tourism industry globally, with the highest number of sites (*n*=101) and annual manta ray focused dives and snorkels (*n* = 157,000) [3]. Japan ranked second, with 145,000 divers visiting only three recognised MRW sites [3].

The Maldives is a hotspot for marine biodiversity [45] and has successfully promoted its natural assets for tourism since 1972, which originated through the focal activity of diving [46]. Tourism visitor arrivals have consistently risen over time, with a 42% increase in just over a decade (2011 = 931,333; 2021 = 1,321,937) [47,48]. Tourists from Asia and Europe were consistently the top visitors to the Maldives, with India (21.1%), Russia (16.8%), Germany (7.2%) and the United Kingdom (4.7%) topping arrivals in 2021 [47]. The Gross Domestic Product (GDP) of the Maldives in 2021 was USD$4.7 billion, with over a quarter of this value being generated through tourism (25.8%; US$1.2 billion) [47]. A separate study reported that the tourism industry also provides employment opportunities for a large segment of the population, with 12% of the nation’s labour force being employed in resorts in 2022 [49]. However, the climate crisis, rising sea levels, ocean acidification, overfishing, pollution and habitat degradation are expected to acutely affect the economy of island nations such as the Maldives [46]. The likely repercussions on the economy are exacerbated within the tourism sector as it relies on the nation’s idyllic natural offerings of picturesque resorts and beaches, and rich biodiversity of marine life [46]. The Maldives is recognised as a world-class diving location, with the abundance of megafauna establishing the country as a popular wildlife watching destination (i.e., whale sharks, reef sharks, sea turtles, rays and cetaceans) [9,22,42,50,51]. Manta ray and shark watching are an especially important component of the dive tourism industry [42,50,52]. Shark fishing previously led to declining shark numbers at many dive sites in the Maldives, reducing shark tourism demand and causing considerable economic losses to the dive-tourism industry [9,41]. Contrary to sharks, manta rays have never been commercially targeted by fisheries in the Maldives [53].

The Maldives 26 atolls support the world’s largest known population of reef manta rays (*Mobula alfredi*), estimated to be 3,500 based on mark recapture analysis of a total of ∼100,000 sightings of 6,200 individuals (G. Stevens, pers. comm.) [54]. The estimated population size of oceanic manta rays (*M. birostris*) which occur in the Maldives (at least for some of the year), based on mark recapture estimates, is likely to be at least one order of magnitude greater than the ∼1,000 individuals [55]. Both populations aggregate at numerous predictable locations, with 48 key aggregation sites (>100 sightings) [56]. Among these, the critical foraging site Hanifaru Marine Protected Area (MPA) in Baa Atoll notably records the highest number of *M. alfredi* sightings [56]. Such consistent aggregation supports an extensive MRW tourism industry. When it was last assessed in 2008, the MRW tourism industry in the Maldives was worth USD$15.5 million per year [3,42], with additional revenue streams associated with this industry not accounted for in the calculations. For example, shark diving in the Maldives generates flow-on socio-economic benefits to the local community and government through employee wages and taxation [52], so this revenue should also be considered for MRW. Findings from the previous MRW economic assessment had positive repercussions for the national protection of manta rays such as the designation of a ray sanctuary [38]. In fact, the Maldives provides a valuable case study for the benefits of tourism within the context of an unfished and relatively ‘pristine’ manta ray population. However, given the growth of both tourism in the Maldives and the global MRW tourism industry [38], the current benefits and the country’s reliance on this source of revenue remains undetermined.

Both *M. birostris* and *M. alfredi* are listed as threatened (Endangered and Vulnerable, respectively) on the IUCN Red List due to their declining populations and conservative life history traits, such as low fecundity, long lifespans, and late age at maturation [57–63]. Indeed, the population recovery capacity of manta rays is among the slowest of all elasmobranchs [58,64], making them particularly vulnerable to over-exploitation in target and bycatch fisheries and other anthropogenic impacts [40,65,66]. Vulnerable species such as manta rays are susceptible to minor negative pressures. Although regulated by the Fisheries and Environmental Protection and Preservation Acts (Act No. 5/87; Act No.4/93), poorly enforced shark and ray tourism in the Maldives can be disruptive to natural behaviours, destructive to habitats, and cause sub-lethal injuries to individuals [38,67,68]. Negative pressure can impact population survival, leading to population declines of up to 99% [65,69–74], and is likely to cause socio-economic impacts in countries that benefit from the MRW tourism industry [3,75]. Therefore, estimating the socio-economic and intrinsic value of these species can help to balance the interests of multiple stakeholders (e.g., fisheries, tourism, coastal development) that influence manta rays or the environment they rely on [38].

Here, we present the first national estimate of the total economic benefits of the MRW tourism industry in the Maldives. We extend previous national economic valuations (i.e., 1997, 2011 and 2013 [3,42,44]) using both surveys and data mining on tour operators, tourists and government to develop a national overview of manta ray tourism and estimate its socio-economic importance in the Maldives. We aim to: (1) quantify the economic benefit to businesses, government, and the community; (2) describe the size, scope, impact and management of the industry; (3) estimate the lifetime value per individual of *M. alfredi* and *M. birostris*; and (4) discuss the intrinsic value of manta rays to the local community. We hope that the findings of this study will highlight the national importance of an industry heavily reliant on flagship species, while also providing valuable information to support ray sanctuaries, and greater investment in MPA management and manta ray conservation.

## Methods

### Study Location

The Republic of Maldives is a small island nation in the central Indian Ocean spanning 870 km in length (north to south) and 128 km across (east to west) at its widest (*3.2°N 73.3°E*) (Fig 1). There are 26 geographical atolls with distinct reef systems [76], grouped into 21 administrative regions (17 atoll regions and four cities – including Malé City, the capital) (Fig 1) [38,77]. The administrative regions organise the country for governance [77]. For this study, we have grouped the data by administrative regions, with the exception of Malé City which is amalgamated with Kaafu, to create more similarly sized areas. Thus, for the purpose of this study, the data is organised into 20 administrative regions that are used hereafter (Fig 1). Of the 1,192 islands in the Maldives, 193 are inhabited community islands (i.e., cities, towns, villages, fishing and farming communities with permanent human habitation), a further 172 are occupied with tourist resorts, and the remainder are protected islands, deserted islands, or are used for agricultural or industry purposes [47]. The population of the Maldives (n=515,114) is centrally located with 41% of residents residing in Malé City (*n*=211,908), with a further 46% on other community islands (*n*=236,911), 3% on industrial islands (*n*=13,831), and 10% on resort islands (*n*=52,482) [78].

**Fig. 1.**
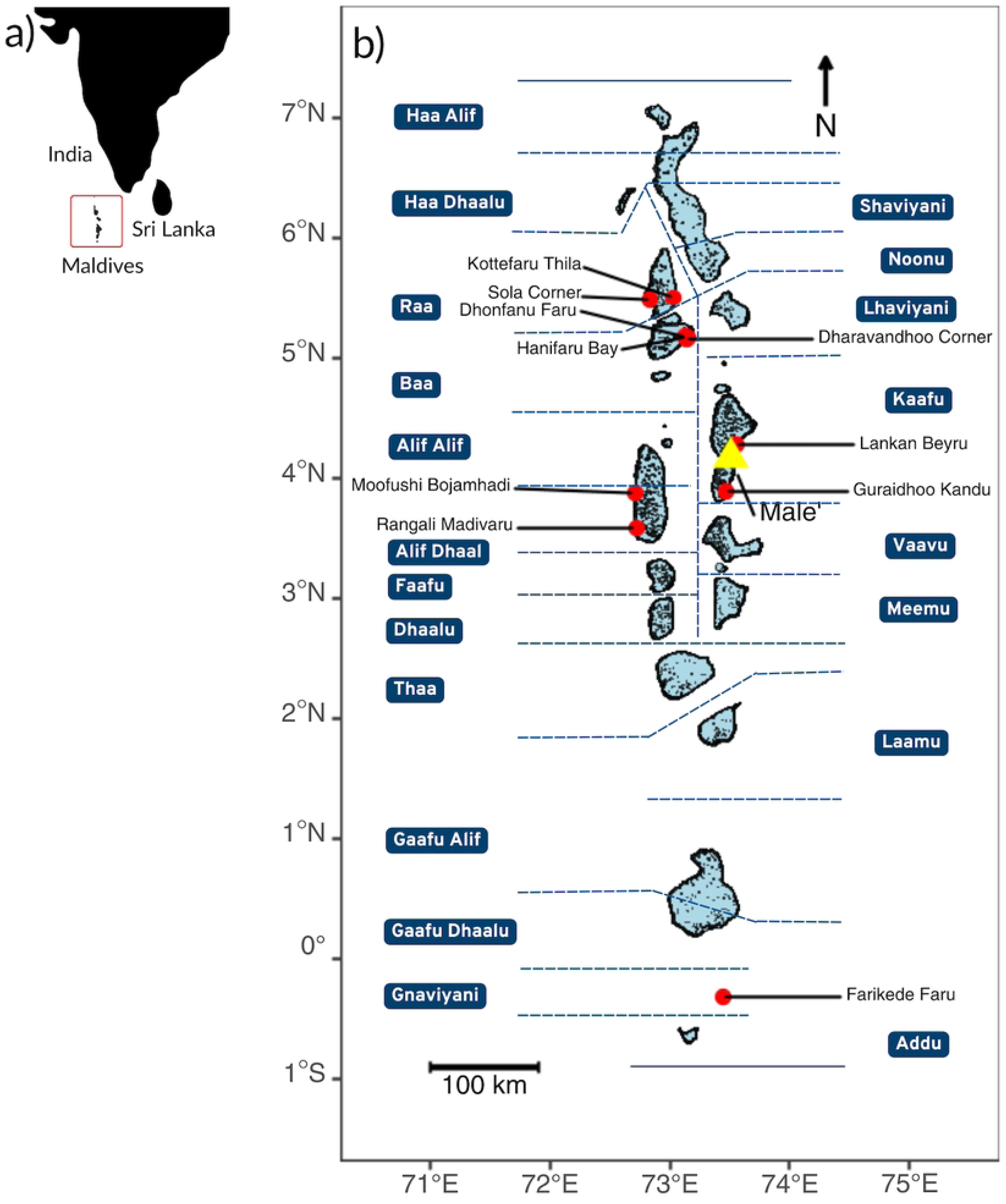
**Geographical distribution of the 10 primary manta ray watching sites (shown by the red circles) in the Republic of Maldives’. The 26 geographical atolls of the Maldives’ archipelago are show by the red box in panel a. The capital, Malé City is identified by the yellow triangle and the nation’s 20 administrative divisions are highlighted in blue in panel b.** Although Malé City is an administrative division, for this study we have grouped it with Kaafu, thus, only 20 divisions are shown in this figure. The manta ray watching sites correspond with those outlined in Table 2.

Tourism is concentrated in the regions surrounding the Velana International Airport (next to Malé City, Kaafu) but increasingly exists across all administrative regions [52,80]. In 2021, there were 354 registered tourism bases, such as resorts, liveaboards and community islands. Within these tourism bases, there were 538 registered tour operators, including dive and excursion activity centres (i.e., excursion centres offer recreational activities including snorkelling trips), and liveaboard vessels [81].

The seasonal South Asian Monsoon strongly influences the oceanography of the Maldives [42]. During the south west (SW) monsoon (May – October), the ocean currents predominantly flow to the east, and in the north east (NE) monsoon (December – March), they flow to the west [42]. The migratory behaviour and the distribution of the Maldives’ manta ray populations are strongly influenced by the monsoon seasons [42,53,56]. *Mobula alfredi* are typically present on the western side of atolls during the NE monsoon and on the eastern side of atolls during the SW monsoon [42,56]. *Mobula birostris* are typically encountered in the south of the Maldives in Gnaviyani (referred to as Fuvahmulah hereafter) between March – April during the transition between the NE and SW monsoons [55].

### Study Design

Data on the revenue of MRW tourism in the Maldives in 2021 were obtained between 2022–2023 from survey responses and online data mining of tour operators and government reports. MRW recreational activities include either snorkelling or scuba diving with the specific intent of viewing *M. birostris* and *M. alfredi* in the wild, in places where manta rays frequent seasonally, periodically, or year-round, consistent with the criteria used in other studies [3,42,56] (see S1 Table for definitions). It can also be watching manta rays from a vessel; however, this has not been quantified in this study. Viewing manta rays that were encountered opportunistically, out of season, or not at a MRW site, were not included as they do not fit the conservative approach of this study as described by Anderson et al. [42]. MRW guests are those who participated in a MRW trip, which may include individuals who participated in multiple trips (S1 Table).

In this study, operators are independently run by third-party companies with separate teams, offices, schedules and vessels. For example, Four Seasons Resort Landaa Giraavaru, a 5-star resort island “base”, has been divided into two “tour operators” as it has both a dive centre and an excursion centre, operated separately. Tour operators are classified into one of the following categories: resorts (*n*=332) that are on private resort islands (*n*=166), local activity centres (*n*=107) that are on community islands (*n*=39), and liveaboards (*n*=99) that are vessels (*n*=99). Thus, tour operators on resorts and community islands are referred to as land-based experiences and liveaboards as boat-based experiences. Because of the stark difference in prices of accommodation and other tourist expenses (*TE*), private resort islands were categorised as either “5-star luxury” (*n*=48) or “regular” (*n*=118) based on the star rating assigned online [82]. Tour operators were further classified according to whether they provided snorkelling or diving activities. Liveaboard cruises offer both snorkelling and diving activities on their trips, but given that most guests onboard are divers, they were considered dive-focused trips and were categorised as a diving centre only.

Private liveaboards are vessels hired privately for multiple-day trips costing between US$12,175 – 84,000, but because of these high prices, it was not possible to decipher the cost per dive and thus cost per MRW dive. Private liveaboards, non-operational or closed bases (*n*=43), were excluded from this study. There were also an additional 613 registered guesthouses [81], however the vast majority did not function as tour operators and outsource snorkelling or diving activities to other businesses on community islands (e.g., dive and excursion centres), and thus were excluded from this study. Hereafter, “tour operators” will include only dive and excursion centres based on resorts, community islands and liveaboard vessels.

#### Tour operator surveys

To obtain information about the diving industry and manta ray tourism for the year 2021, a survey of tour operators was conducted, adapted from O’Malley et al. [3]. Prior to data collection, the survey was first piloted on experts in the industry with feedback incorporated into the survey design. The final online survey was distributed to all tour operators registered with the Ministry of Tourism (n=538) [81] using Google Forms [83] between 26 January – 30 September 2022.

Two survey versions were designed, one for land-based operators (i.e., activity centres in resorts and on community islands) and one for boat-based operators (i.e., liveaboards). The survey consisted of a mix of open-and closed-ended questions, including submitter and tour operator details (questions *n*=11), number of tourists and activity prices for MRW activities (questions *n*=9), manta ray sites and sightings (questions *n*=5), and the socio-economic and intrinsic value of MRW tourism to tour operators, guests and the community (questions *n*=12) (S1 Appendix).

All 538 tour operators in the Maldives were contacted and offered the chance to voluntarily participate in the survey, with 106 operators responding (all providing written consent for their involvement). The almost 20% response rate obtained in our study is comparable to previous studies [52,84]. To estimate the economic benefit of the entire MRW tourism industry in the Maldives and include data on all tour operators, we complemented the survey data with online data mining. Therefore, when registered tour operators did not complete the survey, or when survey data was missing, a standardised assessment was used. This involved data mining the websites and social media platforms of tour operators and travel agencies to collect available MRW tourism data (e.g., activity price, season length and marketing; S2 Table). If there was still missing data for MRW season length, it was sourced from the Manta Trust’s *Maldives Manta Conservation Programme* long-term monitoring database [54] and from Harris et al. [56]. If values could not be sourced, the mean value was calculated by grouping tour operators based on their assigned activity type (i.e., snorkel, dive) and tour operator type (i.e., 5-star resort, resort, local activity centre, liveaboard) from both the survey and mined data. To verify results from survey responses (i.e., activity price and season length), additional data mining was undertaken online for all participating tour operators. Surveys with major discrepancies between operator survey responses and internet data were excluded. For example, if the survey reported an activity price >20% different to that of the same company stated online, the value from the survey was disregarded. Obvious outliers in survey responses were excluded from this study in the initial data cleaning process (*n*=2).

#### Economic benefits of manta ray watching tourism

The total economic benefit from MRW tourism (*TEB_MRW_*, <u>US$</u>; *Equation 1*) in the Maldives in 2021 was estimated as the sum of: i) the tour operator revenue (*TOR*) from manta-ray-specific snorkelling (*TOR_S_*, <u>US$</u>), diving (*TOR_D_*, <u>US$</u>), and diving from a liveaboard (*TOR_LAB_*, <u>US$</u>) activities; ii) the revenue from tourist expenses (*TE*, <u>US$</u>) including accommodation, food and beverages; iii) the revenue for government TAX (*TX*, <u>US$</u>) and staff service charge (*SSC*, <u>US$</u>) (proportions of *TOR+TE*); iv) staff salary revenue (*SSR*, <u>US$</u>) for those who work directly with MRW tourism; and v) the Hanifaru MPA revenue (*MPAR*, <u>US$</u>). Currency values throughout this study are presented as US$.

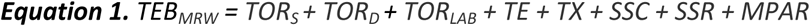

*TOR_S_* and *TOR_D_* (<u>US$</u>) were calculated as the product of the activity price (*AP*, <u>US$</u>), the number of guests per trip (*NG*, <u>unitless</u>), the number of trips per week (*NT*, weeks^-1^), and the number of weeks during which the activities were conducted/season length in 2021 (*NW*, weeks^-1^; *Equation 2a*). Importantly, the revenue generated from tourist visitor entry fees (*EF_V_*, <u>US$</u>) to Hanifaru MPA (*MPAR*) was deducted from *Equation 2a* and remained as revenue generated through MPA management.

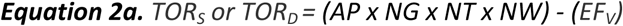

*TOR_LAB_* (<u>US$</u>) was calculated as the product of the value of the activity price (*AP*, <u>US$</u>), the number of opportunities to snorkel and dive with manta rays (*NO*, <u>unitless</u>), the number of guests per trip (*NG*, <u>unitless</u>), and the number of trips per year in 2021 (*NT*, year^-1^; *Equation 2b*). As *TOR_LAB_* already accounted for tourist expenses (*TE*), we estimated that only half of the cost per trip (50%) was used to cover diving expenses, thus used it to calculate the activity price (*AP*). Where values for the activity price (*AP*), the number of guests (*NG*), the number of trips (*NT*), and the number of opportunities (*NO*) were missing from tour operators, a mean value was assigned.

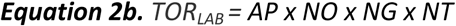

To calculate the combined revenue (*TOR+TE,* <u>US$</u>), which is made up of tour operator revenue (*TOR*) and tourist expenses (*TE*), we determined the base-specific cost benefit transfer ratios using the activity price (*TE:AP,* <u>US$</u>), consistent with other studies [3,22,52,85]. These ratios were calculated by comparing the online price of 10 randomly selected bases (*TE*; including accommodation, food and beverages) with the price of the activity (*AP*; MRW snorkelling or diving). To calculate the tourist expense (*TE*), the mean cost of accommodation per day was used at US$1,733 for 5-star resorts, US$628 for resorts, and US$116 for guesthouses on community islands, combined with a conservative estimate of food and beverage per day at US$150 for 5-star resorts, US$100 for resorts and US$50 on community islands [86]. Given that the price per day on a liveaboard includes food, beverages and accommodation, we assumed that the tourist expenses (*TE*) for this type of operator were 50% of the cost per day, calculated as US$189 for liveaboards. The cost benefit transfer ratios used were 13.265 for 5-star resorts, 7.233 for regular resorts, 2.703 for community islands and 2.586 for liveaboards. For example, if a MRW snorkelling trip cost US$100 (*AP*) at a 5-star resort, there would be an additional US$1,227 in tourist expenses (*TE*) for accommodation, food and beverages, with the combined revenue (*TOR+TE*) calculating to US$1,327. Whereas for the same activity price of US$100 (*AP*) on a community island, there would be an additional US$159 in tourist expenses (*TE*), and the combined revenue (*TOR+TE*) would come to US$259. Given that this data mining exercise was attempted in 2024, three-years after the surveys were distributed, the cost benefit transfer ratios relied on back-calculating the cost of accommodation in 2021 by using the 2024 prices and adjusting for inflation of 12.6% [87]. Tourist expenses are not inclusive of international and domestic travel expenses, despite most of the visitors arriving from overseas via plane and having to travel to their accommodation [47].

The government corporate tax (*TX,* <u>US$</u>) and the staff service charges (*SSC,* <u>US$</u>) in the Maldives in 2021 were 25% of the value of the services provided (i.e., 15% and 10%, respectively) [88]. In this study, the revenue source was calculated for both the tour operator revenue and the tourist expenses (*TOR+TE*). The staff service charge (*SSC*) is generally distributed evenly amongst staff of participating businesses (e.g., hospitality, administration and maintenance staff), and not only to the teams involved in MRW tourism (e.g., dive instructors, boat captains). The staff service charge (*SSC*) is similar to a mandatory gratuity and is paid out to staff on top of the base salary.

MRW employment revenue for 2021 was estimated as the combined value of staff salaries (*SSR*; *Equation 3*) using the number of local Maldivian (*M*) and foreign (*F*) employees (*E*) that work regularly on MRW focused trips including the following positions: snorkel/dive guides, boat crew/captain, administration staff and MPA management staff. The weekly staff wage (*SW*, <u>US$</u>) was conservatively assumed as US$164.3 for Maldivian (*M*) and US$201.2 for foreign (*F*) staff following Zimmerhackel et al. [52]. The annual staff salary was calculated based on the number of weeks per year (*NW,* weeks^-1^) dedicated to MRW activities. When the number of tour operator employees (*M* or *F*) were missing, a mean value based on the tour operator type was assigned

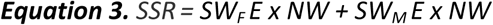

The revenue generated from the management of the Baa Atoll Biosphere Reserve, specifically the Hanifaru MPA (*MPAR*), was used to assess the economic benefits of managing MRW tourism [89]. The revenue generated for management of the Hanifaru MPA (*MPAR*, <u>US$</u>) included resident (*EF_R_*, <u>US$</u>) and tourist visitor (*EF_V_*, <u>US$</u>) entry fees, videography permit fees (*VP*, <u>US$</u>), tour guide licence fees (*TGL,* <u>US$</u>) and professional partnerships fees (*PP*, <u>US$</u>) for streamlined site access (*Equation 4*).

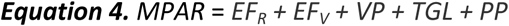

To estimate the revenue generated from tourist visitors at Hanifaru Bay (*REV_HB_*, <u>US$</u>) by tour operators and management; the mean activity price of a snorkel in Baa (*AP_BA_,* <u>US$</u>) and the number of tourist visitor entry fees (*N x EF_V_*, <u>unitless</u>) was used (*Equation 5*). This excluded residents (*R*; Maldivian and work-permit holders) and their entry fees as these are either discounted or not charged by the resorts and activity centres.

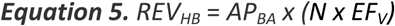

The national contribution of both the tourism [47,48] and fisheries industries [90] were compared to the GDP Market Price of the Maldives. All values used in this study were converted to US$ for consistency (the conversion rate used was Maldivian Rufiyaa (MVR) to United Stated Dollar (US$) on 01/01/2022 at 0.064826) [91].

#### Lifetime value of a manta ray

For both *M. alfredi* and *M. birostris*, the lifetime value (*LV,* <u>US$</u>) of an individual animal for both manta ray species over its lifetime in tourism (*Equation 6*) was calculated using: i) the estimated population size (*P,* <u>unitless</u>); ii) the estimated lifespan (*LS*, <u>years)</u>; iii) the atoll-specific tour operator revenue (*TOR_AS_,* <u>US$</u>) and tourist expenses (*TE_AS_,* <u>US$</u>) for MRW in 2021, consistent with the criteria used in other studies [3,4] (*Equation 6*).

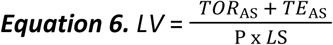

The population of *M. alfredi* was assumed to be 3,500 individuals [54] and for *M. birostris* 1,000 individuals [55]. The estimated lifespan for *M. alfredi* is ∼40 years [92], and because the lifespan for *M. birostris* is unknown, we assumed it to be equal to that of *M. alfredi*. Both *M. alfredi* and *M. birostris* frequent Maldivian waters, but there is only one region where *M. birostris* frequents reliably, Fuvahmulah. To calculate the lifetime value (*LV*) for the two species, it was necessary to split the combined revenue into atoll-specific (*TOR_AS_ + TE_AS_*) visitation for bases that are moveable (i.e., liveaboards) and for bases that contain permanent structures (i.e., resorts, community islands). This is because liveaboards visit multiple regions during a single trip, while tour operators with permanent structures generally operate within a single region. For *M. birostris*, the revenue from the tour operators was generated in Fuvahmulah only, while the revenue for *M. alfredi* corresponds to all the other regions they are sighted seasonally. We estimated that 4% of the revenue generated (*TOR_AS_ + TE_AS_*) from liveaboards was from time spent in Fuvahmulah and the other 96% of revenue was from the remaining regions, based on the responses on visitation frequency from eight tour operators.

#### Intrinsic value of manta rays

The importance of manta rays for business operations, local communities, and education outreach were evaluated by quantifying the frequency of the words used to answer three open-ended questions in the tour operator surveys, including: 1. Why/why-not do you consider manta rays to be important to your business; 2. Why/why-not do you consider manta rays to be important to local communities in the Maldives; and 3. Do you think that your manta ray trips helped to educate guests about manta rays and conservation, and how? These data were processed through systemising descriptive responses to identify patterns and themes behind textual data. This data was first visualised using wordclouds created in R (version 4.4.1) [93]. Text mining and removal of stop words was done using the ‘tidytext’ R package [94]. Some of the words used in the questions were also removed pre-analysis e.g., “manta”, “mantas”, “manta ray”, “Maldives”. Additionally, eight visitor surveys conducted by the Ministry of Tourism between 2011—2021 (visitor participants *n*=11,625) were used to assess the primary motivations of tourists, visitor satisfaction and diving/snorkelling participation [95].

## Results

### Scale of manta ray watching tourism

A total of 80% (*n*=282) of the 354 tourism bases in the Maldives offered MRW diving and/or snorkelling in 2021 (Table 1). Within these tourism bases, 69% (*n*=374) of the 538 registered tour operators participated in MRW tourism. This industry is widely distributed across the Maldives, being present in all but one of the 20 administrative regions used in this study. However, land-based experiences were concentrated primarily in Kaafu, Alif Dhaal and Baa, which had 76, 46, and 43 tour operators, respectively. Whereas liveaboard experiences generally visited multiple regions, with Alif (both Alif Alif and Alif Dhaal), Kaafu and Meemu being the most visited during the NE Monsoon, and Baa, Kaafu and Alif in the SW Monsoon.

**Table 1.** Registered, verified, and operational tour operators in the Republic of Maldives (established by the Ministry of Tourism and verified by this study; *n*=538) and the extent of manta ray watching (MRW) tourism.

| Tour operator type | Base type [$n$] | Tour operators [$n$] | Tour operators offering MRW [$n$ (%)] | Tour operators (separated into activity centre type) offering MRW [$n$ (%)] | | Tour operators that completed the survey [$n$ (%)] | Survey participants – offering MRW [$n$ (YES)] |
| --- | --- | --- | --- | --- | --- | --- | --- |
|  |  |  |  | Dive Centre | Excursions Centre |  |  |
| 5-star luxury resorts | Private resort island ($n=48$) | 96 | 62 (65%) | 35 (73%) | 27 (56%) | 21 (22%) | 20 |
| Regular resorts | Private resort island ($n=118$) | 236 | 139 (59%) | 80 (68%) | 59 (50%) | 43 (18%) | 42 |
| Local centres | Inhabited community island ($n=40$) | 107 | 88 (82%) | 66 (80%) | 16 (94%) | 29 (27%) | 27 |
| Liveaboards | Vessel ($n=99$) | 99 | 85 (86%) | 85 (86%) | | 13 (13%) | 12 |
| **Total** | **354** | **538** | **374 (70%)** |  |  | **106 (20%)** | **101** |

Tour operators reported 92 MRW sites for diving/snorkelling with manta rays. Within these sites, 65 were regularly visited for diving and 56 for snorkelling (some sites were used for both activities). They identified Hanifaru MPA in Baa (*n*=18) and Lankan Beyru in Kaafu (*n*=17) as the two primary MRW sites (Table 2). Hanifaru MPA was the most important MRW site in terms of number of tourists per trip (*n*=52.6 ± 6.1 Standard Error (SE)), number of boats per trip (*n*=8.8 ± 1.2 SE), and the number of manta rays sighted per trip (*n*=32.3 ± 4.0 SE). Most of the MRW sites were primarily frequented by *M. alfredi.* Encounters with *M. birostris* were only promoted in Fuvahmulah.

**Table 2.** Most visited manta ray watching (MRW) dive and snorkel sites as ranked by tour operators listed in their top five, and the mean (± *SE*) number of tourists, tourist vessels and manta rays per trip. The active MRW season is categorised as either the northeast (NE) or southwest (SW) monsoon.

| Region | Manta ray watching site name | Latitude (°) | Longitude (°) | Species primarily encountered | Number of tour operators that listed this site [$n$] | Monsoon active | Activity | Mean no. tourists per trip [$n \pm SE$] | Mean no. manta rays per trip [$n \pm SE$] | Mean no. boats per trip [$n \pm SE$] |
| --- | --- | --- | --- | --- | --- | --- | --- | --- | --- | --- |
| Baa | Hanifaru Bay | 5.17 | 73.15 | *M. alfredi* | 18 | SW | Snorkelling | 52.6 (± 6.1) | 32.3 (± 4.0) | 8.8 (± 1.2) |
| Kaafu | Lankan Beyru | 4.28 | 73.56 | *M. alfredi* | 17 | SW | Both | 26.0 (± 4.5) | 4.1 (± 0.4) | 5.1 (± 0.8) |
| Raa | Sola Corner | 5.49 | 72.83 | *M. alfredi* | 12 | SW | Both | 15.3 (± 4.5) | 4.0 (± 0.3) | 3.1 (± 0.3) |
| Alif Dhaal | Rangali Madivaru | 3.59 | 72.72 | *M. alfredi* | 12 | NE | Both | 32.4 (± 7.3) | 4.8 (± 0.4) | 6.8 (± 1.3) |
| Baa | Dhonfanu Faru | 5.18 | 73.12 | *M. alfredi* | 11 | SW | Both | 22.5 (± 7.5) | 11.4 (± 4.9) | 4.4 (± 1.1) |
| Raa | Kottefaru Thila | 5.51 | 73.03 | *M. alfredi* | 11 | NE | Both | 10.0 (± 2.1) | 6.3 (± 1.0) | 2.8 (± 0.3) |
| Alif Dhaal | Moofushi Bojamhadi | 3.88 | 72.71 | *M. alfredi* | 11 | NE | Both | 35.8 (± 5.9) | 4.6 (± 0.7) | 3.4 (± 0.4) |
| Baa | Dharavandhoo Corner | 5.16 | 73.14 | *M. alfredi* | 10 | SW | Both | 13.3 (± 0.9) | 3.0 (± 0.0) | 3.5 (± 0.2) |
| Fuvahmulah | Farikede Faru | -0.32 | 73.45 | *M. birostris* | 10 | NE | Diving | 15.7 (± 6.7) | 3.8 (± 1.4) | 2.5 (± 0.6) |
| Kaafu | Guraidhoo Kandu | 3.89 | 73.47 | *M. alfredi* | 10 | SW | Diving | 13.5 (± 2.9) | 3.3 (± 0.4) | 2.3 (± 0.5) |

### Economic benefits from manta ray watching tourism

Tour operator revenue (*TOR*) for manta-ray-specific snorkelling, diving, and diving on a liveaboard in the Maldives in 2021 was estimated to be US$39,027,696 (Table 3) from 475,061 guests participating in MRW experiences (diving *n*=350,689, snorkelling *n*=124,372). Most of this revenue is generated from MRW experiences in three regions, including Kaafu, Alif Dhaal and Baa, which generate approximately 12% (US$4.8 million), 15% (US$5.8 million), and 18% (US$6.9 million) of the total tour operator revenue (*TOR*) respectively (Fig 2). Five-star resorts contributed the least to the tour operator revenue when compared with other tour operator types (Fig 3e). Despite having the highest activity prices (*AP*), 5-star resorts had the lowest number of guests per trip (*NG*), trips per week (*NT*), and season length (*NW*; Fig 3a, b, c, d). The mean activity price (*AP*) across operators was US$86.6 ± 2.3 for diving and US$121.8 ± 6.6 for snorkelling. The perceived value to businesses of an opportunistic encounter with a manta ray was high, as demonstrated by 81% of tour operators that featured manta rays in their operation’s marketing (*n*=448; i.e., logo, photos on website, marketing mentions famous manta ray dive sites). Benefits from MRW tourism also reach other local businesses. Tourist expenses (*TE*) from MRW dive/snorkel guests were an estimated US$188,270,681 for local businesses, including accommodation, food and beverages (Table 3). The combined revenue (*TOR+TE*) of MRW was estimated at US$227,298,377 and contributed an estimated 4.8% to the annual 2021 GDP Market Price in the Maldives (Table 4). Tourism constitutes a significant portion of the Maldivian economy, contributing over 25% to its industry to its GDP in 2021 [47].

**Fig 2.**
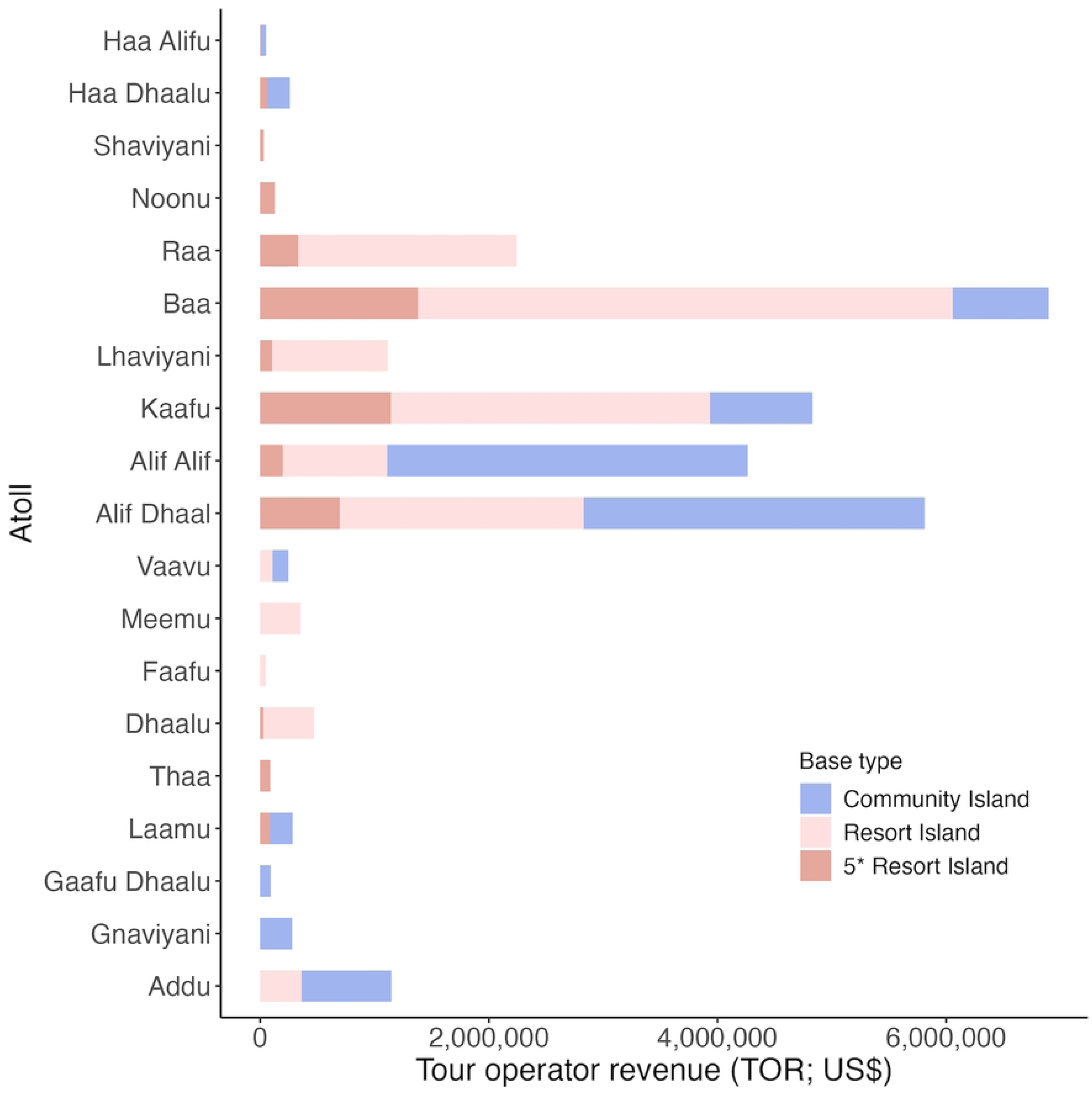
**Estimated tour operator revenue (*TOR*) and distribution of manta ray watching tourism in the Republic of Maldives across tourism bases** (all values in US$). This excludes liveaboards as they generally visit multiple regions during a single trip. Each bar represents a single administrative region (*n*=19). Regions are ordered from north (Haa Alif) to south (Addu). Malé City is grouped with Kaafu. There is no reported manta ray watching tourism in Gaafu Alifu, thus it did not generate revenue and it is not displayed on this figure.

**Fig 3.**
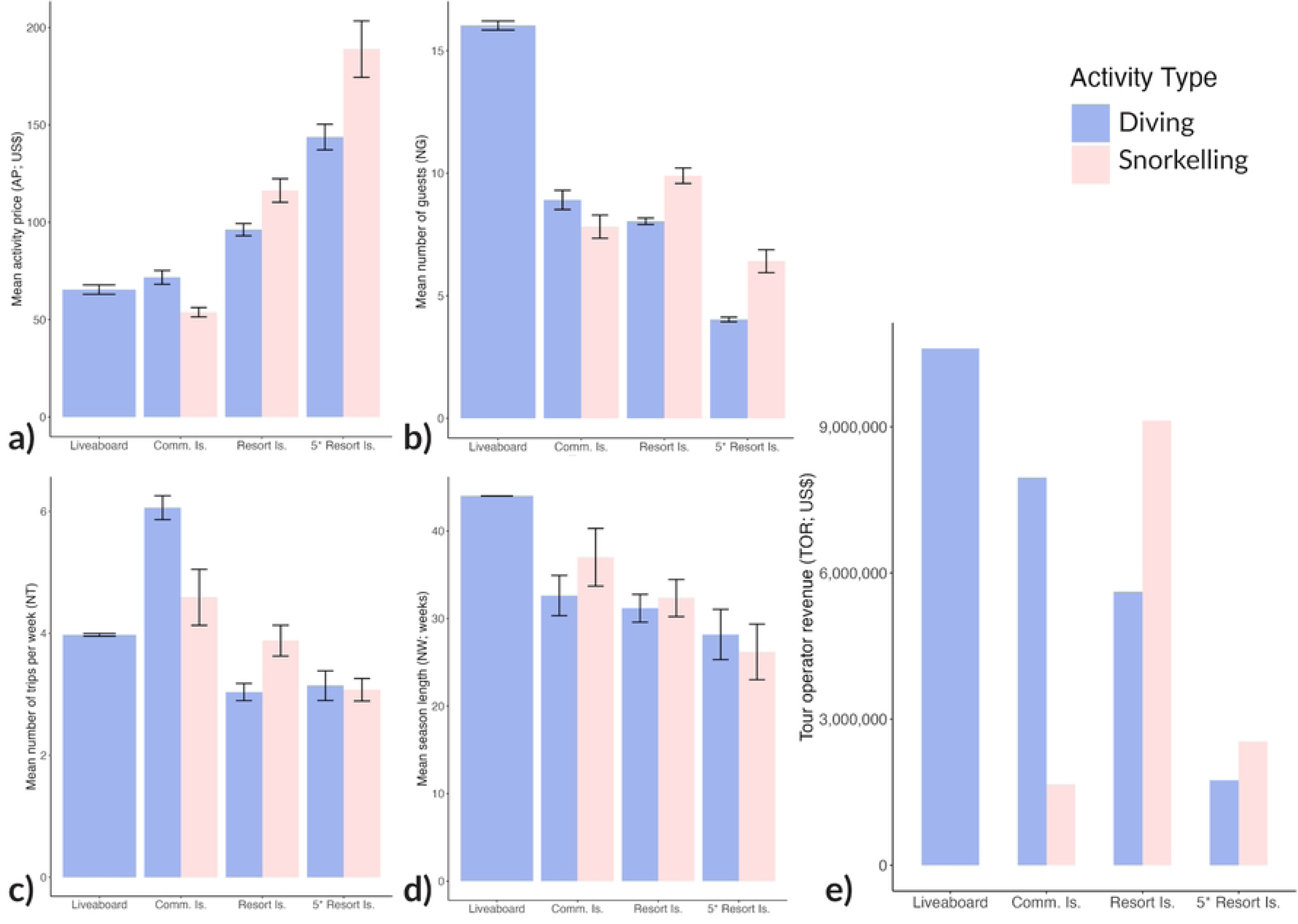
Tour operator revenue (*TOR*) generated through manta ray snorkelling and diving activities categorised by tour operator type in the Republic of Maldives in 2021. This was calculated as a product of a) activity price per guests (*AP*), b) number of guests per trip (*NG*), c) number of trips per week (*NT*) and d) season length (in weeks) per year (*NW*), displayed here as the Mean (± *SE*) value (All values in US$). Comm. Is., community Island; Resort Is., resort island; 5*Resort Is., 5-star resort island.

**Table 3.** Economic benefits of manta ray watching (MRW) tourism in the Republic of Maldives in 2021 collected via internet research and tour operator surveys. Tourist expenses (*TE*) are additional costs and are not included in the tour operator revenue (*TOR*). The 374 tour operators that offer MRW tourism are categorised based on the activity type they offer (diving-specific operators *n*=267 and snorkelling-specific operators *n*=107). Tour operators sold 475,061 tickets to take guests on a MRW trip (diving-specific trips *n*=350,689 and snorkelling-specific trips *n*=124,372) (All values in US$).

| | Revenue source | Diving-specific revenue (*D*) (US\$) | Snorkelling-specific revenue (*S*) (US\$) | Total (US\$) |
| --- | --- | --- | --- | --- |
| *TOR<sub>D/S/LAB</sub>* | Tour operator revenue | \$25,929,181 | \$13,098,515* | \$39,027,696 |
| *TE* | Tourist expenses | \$97,418,749 | \$90,851,933 | \$188,270,681 |
| *TX* | Tax | \$18,502,189 | \$15,592,567 | \$34,094,757 |
| *SSC* | Staff service charge | \$12,334,793 | \$10,395,045 | \$22,729,838 |
| *SSR* | Employee staff salaries | \$21,636,162 | \$5,352,606 | \$26,988,768 |
| | **Sub total** | \$175,821,074 | \$135,290,665 | \$311,111,739 |
| *MPAR* | Management | | \$252,891 | \$252,891 |
| *TEB<sub>MRW</sub>* | **TOTAL economic benefit** | \$175,821,074 | \$135,543,556 | \$311,364,630 |

\*The Hanifaru Bay MPA ticket revenue was deducted from tour operator revenue as this is accounted for by the MPA management (*MPAR*).

**Table 4.** The Gross Domestic Product (GDP) in the Republic of Maldives and the share contribution of national industries including manta ray watching (MRW) tourism for 2008 – 2010, and 2019 – 2021. The GDP and Tourism Contribution values were converted to US$ for consistency (Maldivian Rufiyaa -MVR to United Stated Dollar – US$ on 01/01/2022 @0.064826).

| Year | Number of tourists (n) | GDP Market price (US\$)<sup>a</sup> | Tourism Contribution (US\$)<sup>a</sup> | Share of Tourism Contribution (%)<sup>a</sup> | Fisheries Contribution (US\$)<sup>b</sup> | Share of Fisheries Contribution (%)<sup>b</sup> | MRW tour operator revenue Contribution (US\$) | Share of MRW tour operator revenue Contribution (%) | MRW combined revenue Contribution (US\$) | Share of Combined economic revenue Contribution (%) |
| --- | --- | --- | --- | --- | --- | --- | --- | --- | --- | --- |
| 2008 | 638,012 | \$1,200,983,149 | \$315,707,003 | 26.3 | | | \$8,100,000<sup>c</sup> | 0.7 | \$15,471,000<sup>d</sup> | 1.3 |
| 2009 | 655,852 | \$1,144,065,131 | \$298,722,355 | 26.1 | | | | | | |
| 2010 | 791,917 | \$1,359,160,785 | \$345,877,442 | 25.4 | | | | | | |
| 2019 | 1,702,887 | \$5,007,100,102 | \$1,305,678,593 | 26.1 | \$152,122,804 | 3 | | | | |
| 2020 | 555,494 | \$3,330,093,026 | \$483,284,540 | 14.5 | \$167,843,327 | 5 | | | | |
| 2021 | 1,321,937 | \$4,720,241,070 | \$1,217,838,143 | 25.8 | \$163,804,611 | 3.5 | \$63,108,424 | 0.8 | \$227,298,377 | 4.8 |

Socio-economic benefits of MRW tourism flow into the community in the form of staff salary revenue (*SSR*). The tour operator surveys revealed that operators employ 3,876 staff (Maldivian *n*=2,602, Foreign *n*=1,262) who regularly work on MRW trips (Table 5). In addition, 12 Maldivians were employed to manage the Baa Atoll Biosphere Reserve [89]. The mean annual salary for Maldivian staff (US$8,544) was 81.7% that of foreign staff (US$10,464). Calculated only for the number of weeks (*NW*) of the MRW season, the total staff salary revenue (*SSR*) for tour operator staff was estimated at US$26,886,240 (Maldivian = US$14,675,591, Foreign = US$12,210,649) and for the Biosphere Reserve staff (Maldivian = US$102,529). Local Maldivian staff make up 67% of the workforce employed to work in MRW tourism and management and receive 55% of the distributed staff salary revenue (*SSR*). The flow-on benefits of this industry additionally extend to the government through tax (*TX*; US$34,094,757) and then again to the workforce (not only those employed in roles to directly work in MRW tourism) in the form of staff service charge (*SSC*; US$22,729,838; Table 3).

**Table 5.** Staff employed in operations that actively participate in manta ray watching tourism. . Calculations only account for respective manta ray watching seasons. *Weekly salaries were obtained from Zimmerhackel et al. [52].

| Code | Salaries | Foreign Staff<br>[*n* (*F*)] | Maldivian Staff<br>[*n* (*M*)] | Total Staff<br>(*n*) |
| --- | --- | --- | --- | --- |
| *E* | Number of employees working with MRW diving | 1,024 | 1,893 | 2,781 |
| *E* | Number of employees working with MRW snorkelling | 238 | 709 | 1,083 |
| *E* | Number of employees working in Management | 0 | 12 | 12 |
|  | **Total employees** | 1,262 | 2,614 | 3,876 |
| *SW* | Weekly salaries (US\$) | **\$201.23\*** | **\$164.31\*** | |
| ***SSR*** | **Total staff salary revenue (US\$)** | \$12,210,649 | \$14,778,119 | \$26,988,768 |

In terms of management, the single monitored and ticketed MPA in the Maldives, Hanifaru, in the Baa Atoll Biosphere Reserve (*MPAR*), generated US$252,891 in revenue for management bodies in 2021 (Table 6). This includes the visitor (*EF_V_*) and resident entry fees (*EF_R_*), videography permit fees (*VP*), tour guide licence fees (*TGL*) and professional partnerships (*PP*). This MPA was the top visited MRW site in the Maldives, as indicated by tour operator surveys (Table 2). On average, tour operators charged US$137.3 ± 14.3 per snorkel trip (*AP*) in Baa. The 11,258 visitors to Hanifaru MPA generated an estimated US$1,319,171 in revenue for tour operators and management (*REV_HB_*). The total economic benefits (*TEB_MRW_*) of MRW tourism in the Maldives in 2021 was estimated at US$311,364,630 (Table 3).

**Table 6.**
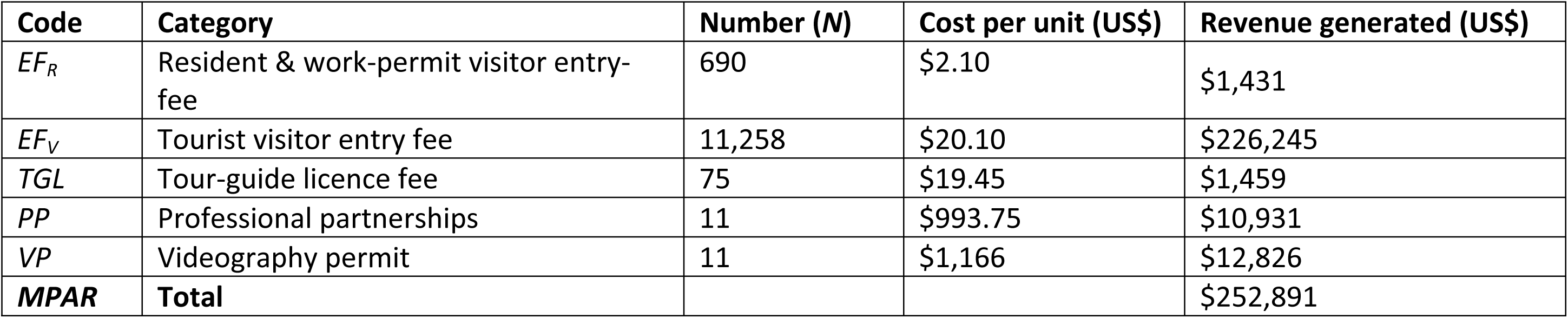
Revenue generated for management bodies from Hanifaru Bay Marine Protected Area (*MPAR*). Amount (*N*) and cost per unit sourced from Baa Atoll Biosphere Reserve [89].

| Code | Category | Number ($N$) | Cost per unit (US\$) | Revenue generated (US\$) |
| --- | --- | --- | --- | --- |
| $EF_R$ | Resident & work-permit visitor entry-fee | 690 | \$2.10 | \$1,431 |
| $EF_V$ | Tourist visitor entry fee | 11,258 | \$20.10 | \$226,245 |
| $TGL$ | Tour-guide licence fee | 75 | \$19.45 | \$1,459 |
| $PP$ | Professional partnerships | 11 | \$993.75 | \$10,931 |
| $VP$ | Videography permit | 11 | \$1,166 | \$12,826 |
| **MPAR** | **Total** | | | \$252,891 |

### Lifetime value of a manta ray

The lifetime values (*LV*) were calculated using the atoll-specific generated revenue; for *M. birostris* using revenue from Fuvahmulah (*TOR_AS_* = US$705,036 and *TE_AS_* = US$1,575,327), and for *M. alfredi* using the revenue from the other regions (*TOR_AS_* = US$38,322,660 and *TE_AS_* = US$186,695,354). An individual *M. alfredi* was estimated to be worth US$2,571,634, compared to a value of US$91,215 for *M. birostris* during their lifetime (Fig 4).

**Fig 4.**
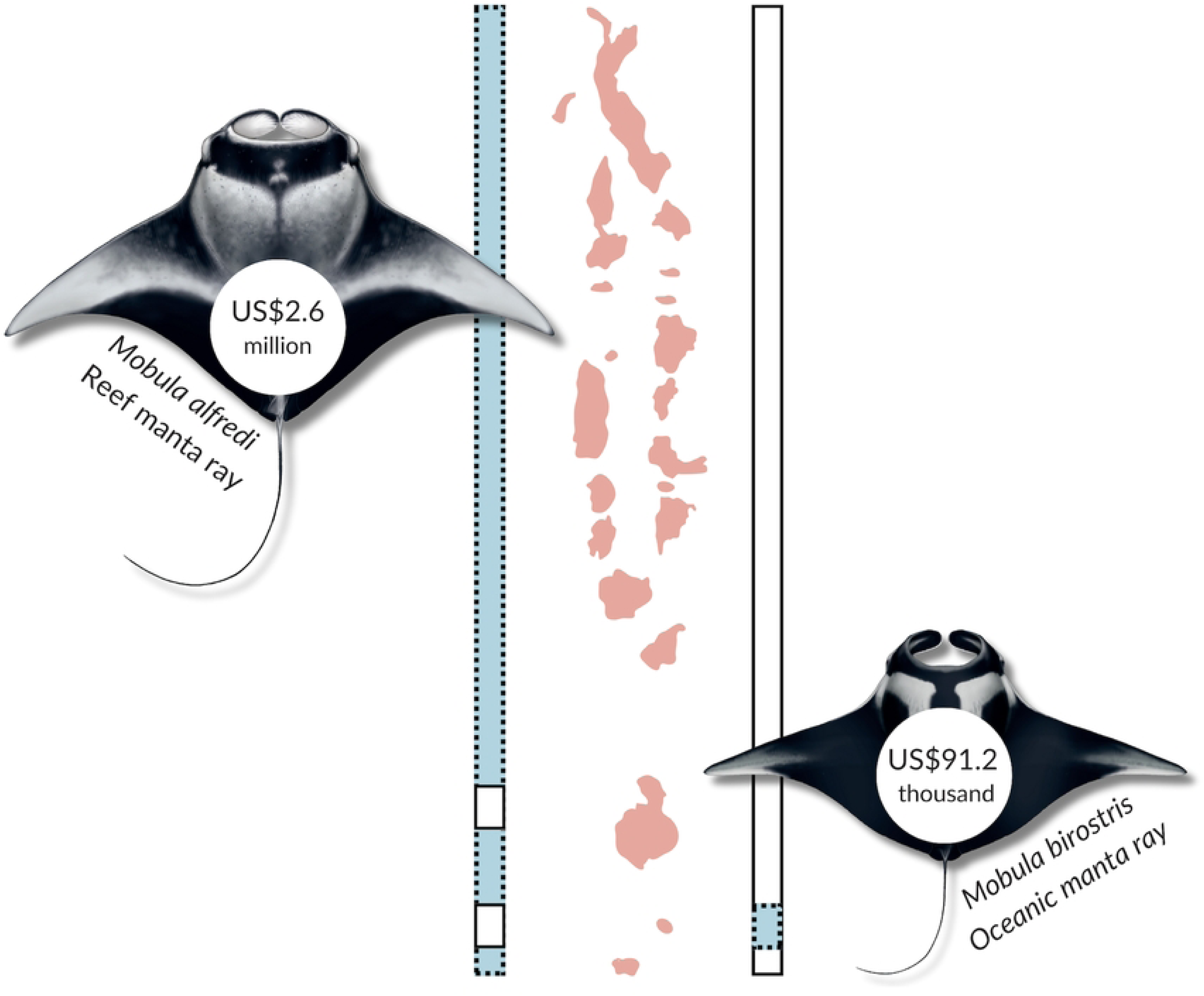
The lifetime value of individual manta rays (both species) as part of tourism calculated using the combined economic revenue from manta ray watching in the Republic of Maldives for 2021. Tourism that focused on *Mobula alfredi* (reef manta rays) was identified in 18 administrative regions, compared to only one region for *M. birostris* (oceanic manta rays; Gnaviyani/Fuvahmulah). The geographical extent of manta ray tourism is shown with the coloured dotted rectangle for both species.

### Intrinsic value

Tour operator survey respondents provided insight into the intrinsic value of manta rays to their businesses, to in-country visitors, and to local communities (Fig 5). All tour operators ranked manta rays in the top five sea life that visitors most wanted to see (*n*=100) with 75% ranking them in their top two, and 28% ranking them as the top attraction. Operators perceived manta rays to be important to their business (yes = 97%, *n*=101) and to the wider local community (yes = 99%, *n*=101; Fig 5). The word count analysis showed the most frequently used words by survey responders to describe why they believed manta rays to be important to their operation (n=103) were: “guests” (*n*=28), “people” (*n*=18), “season” (*n*=10), “divers” (*n*=8), resort” (*n*=8) and “trips” (*n*=8; Fig 6a). It was also reported that manta rays in the Maldives were a “bucket list” item for many guests (*n*=9). Compared to when survey respondents were asked to describe the importance of manta rays to the community (*n*=102), the most frequently used words were: “local” (*n*=25), “tourism” (*n*=24), “income” (*n*=16), “communities” (*n*=15) and “tourists” (*n*=12; Fig 6b). One survey respondent also reported that “Mantas have a certain cultural significance to local communities in the Maldives as well as attracting tourists to the area along with the associated economic benefits”.

**Fig 5.**
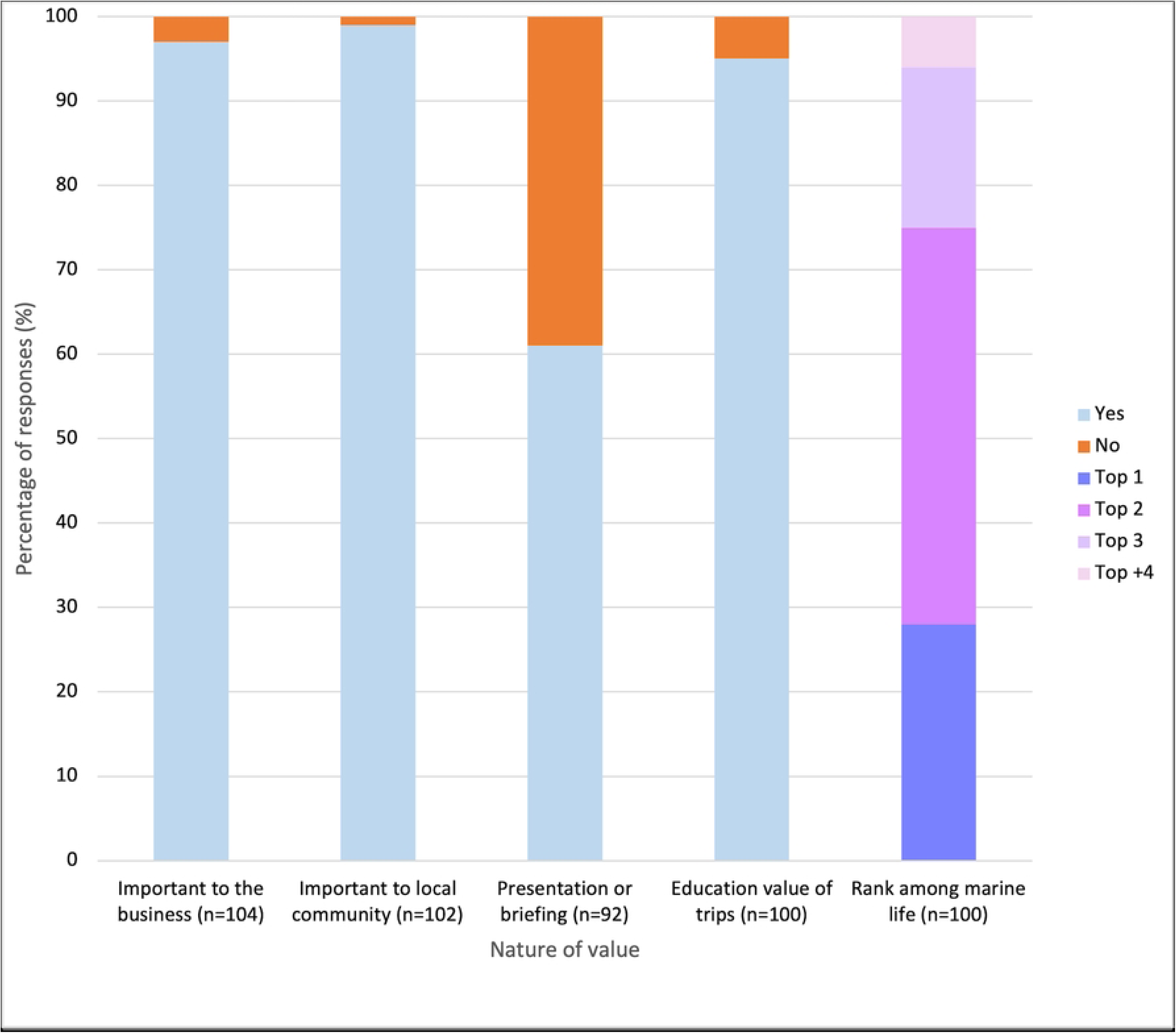
The perceived socio-economic importance of manta ray watching tourism to tour operators (proportion) in the Republic of Maldives. These responses were via survey questions about the nature of the value e.g., to their business, the community, as a tool for education and conservation for guests, and how manta rays ranked among marine life visitors wanted to see.

**Fig. 6.**
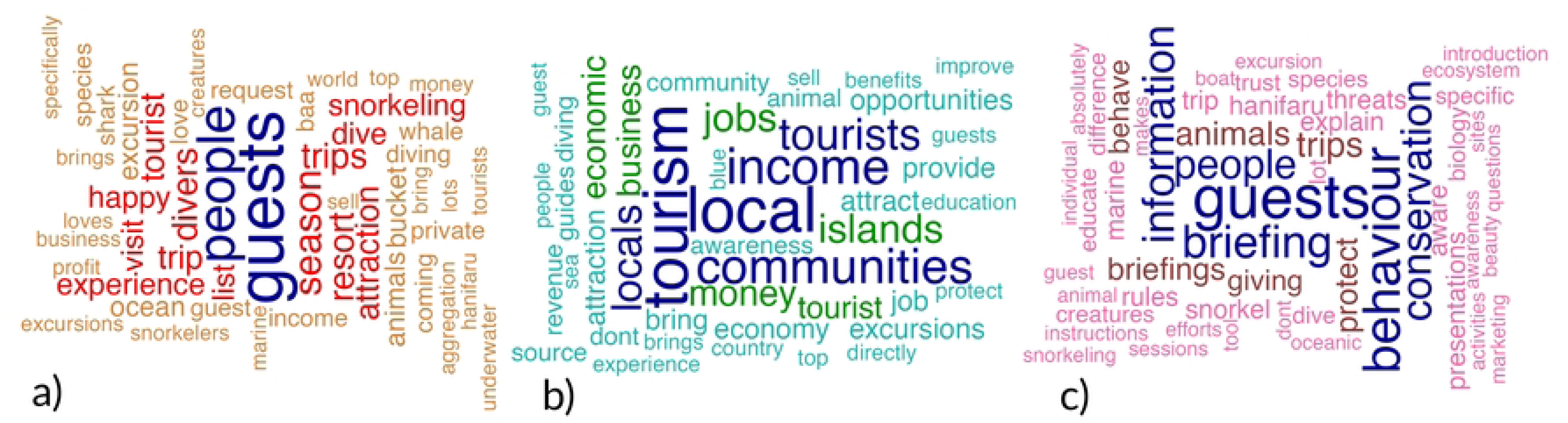
**Most common words used by tour operators in the Republic of Maldives to describe the importance of manta ray watching (MRW) tourism to a) their operation (*n*=103), b) to the community (*n*=102) and c) as a tool for education and conservation for guests (*n*=100) visualised in wordclouds.** The wordclouds show how many times individual words (max words = 70) were detected in their statements. This corresponds to the size and colour of the words.

The value of manta rays was additionally demonstrated with 61% of survey respondents (*n*=56) reportedly offering an educational presentation and/or briefing to trip guests through their operation (Fig 5). These presentations generally covered aspects of biology, threats to and conservation of manta rays, and how to swim responsibly with these animals. Without taking into consideration tour operators that did not complete the survey, it is estimated that at least 17,655 MRW guests received a presentation or briefing as part of their MRW trip experience in 2021. Most tour operators believed that taking guests to see manta rays helped to educate and conserve these animals (95%, *n*=95). The most frequently used words by survey respondents to describe why they believed that manta ray trips helped to educate guests about manta rays and conservation (*n*=100) were: “guests” (*n*=26), “behaviour” (*n*=19), “briefing” (*n*=15), “information” (*n*=14), and “conservation” (*n*=13; Fig 6c).

## Discussion

### Economic benefits of manta ray watching tourism

We show that marine wildlife tourism can offer immense value to local and national economies by generating revenue, supporting cultural heritage, and creating employment opportunities. Using the Maldives as a case study, this study provides an updated national evaluation of the economic benefits of MRW tourism. We found that the 475,000 dive and snorkel trips taken by MRW guests in the Maldives generated an estimated US$227.3 million (US$39 million in revenue for tour operators and US$188.3 million in tourist expenditures) for the local tourism industry in 2021. These estimates are a substantial increase from previous assessments of MRW tourism in the Maldives in 1997 (US$7.8 million) and 2008 (US$8.1 million) [41,42]. When the industry was last valued it was estimated to generate a total of US$15.5 million in combined revenue, including US$7.4 million in tourist expenditures from the 157,000 MRW trips [3]. The combined revenue provides a conservative baseline for the direct economic impact of wildlife tourism [2–4,22,24,25,96,97]. Although differences in methodologies by the studies means our estimate is not precisely comparable to the 2008 estimate, the 2021 value reflects a remarkable >380% growth in tour operator revenue over the past decade. This growth underscores the increasing economic significance of MRW tourism. Additionally, it mirrors that of the overall Maldives’ tourism sector which reported 286% growth in the tourism contribution to GDP over the same period (2008 = US$515.7 million and 2021 = US$1.2 billion) [47].

Alongside the growth in revenue from tourism, the annual international visitor numbers to the Maldives have doubled (2008 = 638,012 and 2021 = 1,321,937) [47]. During routine departure surveys from 2009 – 2022 (*n*=11,625), over half (55%) of the international visitors cited the “underwater beauty” of the marine environment as the primary reason for their in-country visit [95]. Many international visitors were also specifically attracted by diving and snorkelling (26%), and particularly by megafauna, including manta rays (22%) [95]. Reef sharks and whale sharks (*Rhincodon typus*) are also economically important elasmobranch megafauna in the Maldives, with both industries reporting revenue growth over recent years [50,52]. Although mindful of methodological differences, reef shark diving contributed US$14.4 million in tour operator revenue in 2016, an 83% increase from US$7.9 million in 1992 [41,52]. While whale shark watching contributed US$9.4 million in 2013, rising from US$7.6 million in 2012 [50], the industry therefore growing 24% in one year (no difference in methods). Although reef and whale shark tourism in the Maldives show industry specific growth, they do not demonstrate growth of a similar magnitude to MRW tourism.

Tour operators ranked manta rays in the top five sea life that diving and snorkelling tourists wanted to see, with 28% ranking them as the top attraction. Manta rays also generate value through marketing and are used as a selling point for businesses to attract potential customers. Despite some operators not directly selling manta-ray-focused trips, the perceived value of even an opportunistic encounter holds substantial business appeal, highlighting the profitability of promoting this kind of tourism. Consistent with the increased number of international visitors, the Maldives reported a 272% increase in registered bases (including guesthouses) from 250 in 2008, to 929 in 2021 [47,48]. Moreover, during the same timeframe, the mean price of a MRW diving trip nearly doubled, from US$45–70 to US$87, and snorkelling trips saw a six-fold increase from US$20 to US$121.8 [42]. The large increase in the price for a MRW snorkelling trip could be attributed to the rise in popularity of this kind of activity, particularly at resort bases which are generally more expensive [86].

MRW tourism contributed substantially to the Maldives’ economy, representing 4.8% of its GDP in 2021, surpassing that of the fishing and fish processing industry, which contributed 3.5% to the GDP in the same year [90]. The Maldives is a tuna fishing nation [98] and has an especially high dependence on the sustainable availability of marine resources for both food security and economic prosperity [35]. As a country that does not harvest its manta rays [38], but is reliant on both fisheries and tourism industries, the Maldives provides an example of how these two industries can co-exist when managed sustainably. This contrasts with manta ray tourism in Mozambique, which reported a 92.5 to 99% decrease in manta ray sightings over the 20-year study period in one region of the country where MRW is an important part of the local economy [73]. Fisheries mortality was identified as having played a significant role in this decline [73].

The contribution of MRW to the Maldives economy exceeds estimates from older studies in other nations that used similar survey design and analysis [3,8,42]. Nevertheless, the comparison between Mozambique in 2014 (US$34 million; [8]), the global industry in 2012 (US$140.7 million; [3]), and the Maldives in 2021 (US$227.3 million) demonstrates the huge size of the Maldives industry. In a global review of MRW tourism by O’Malley et al. [3], the Maldives had the highest number of both MRW sites and guests. Given the current estimate for 2021, it is likely that the Maldives remains one of the highest, if not the highest, MRW revenue generating nations world-wide.

Both the Maldivian economy and the livelihood of its people are largely dependent on marine resources [99]. This study provides evidence for the flow-on socio-economic impacts of manta ray tourism in the Maldives. Almost 4,000 staff were employed by tour operators to work directly on MRW diving and snorkelling tourism activities, generating US$27 million per year in staff salary revenue. Despite the 2,602 Maldivian staff representing 67% of the MRW tourism workforce, they only received 55% of the distributed staff salary of all employed within this industry. Similarly, reef shark-diving tourism in 2016 in the Maldives employed 131 Maldivians who represented 55% of the shark-diving workforce but received 50% of the distributed staff salary [52]. The observed discrepancy in staff salary revenue between Maldivian and expatriate foreign staff in both studies likely stems from the disproportionate employment of Maldivian nationals in lower-paid jobs such as divemaster, equipment maintenance, or boat operations [100]. A shortage of trained local personnel often resulting in expatriates occupying managerial and senior positions while locals are largely confined to less skilled, lower-paid jobs is a pattern recognised in many Global South tourism industries [101]. While some operators in the Maldives are already implementing initiatives to further address these employment disparities and maximise local economic benefits, encouraging the widespread adoption of a structured approach could be beneficial. Such an approach would prompt businesses not only to prioritise local employment but also to invest in upskilling opportunities, thereby preparing Maldivian staff for promotions and managerial roles. Beyond fostering individual career progression, this model has been shown to offer several broader benefits, including enhanced local economic growth, improved skill development within the community, long-term economic sustainability and reduced recruitment costs [100,102]. For the full economic benefits of tourism to be realised and retained locally, nurturing local capacity is indispensable [102].

Socio-economic benefits also flow-on to reach the government via tax and MPA management, and to staff employed in the hospitality and tourism industries through staff service charge. These diverse revenue streams underscore the industry’s substantial local importance. In 2021, the total economic and socio-economic benefits of MRW tourism to businesses, government and the community in the Maldives, considering both direct and indirect revenues, exceeded US$311 million. While detailed economic valuations of wildlife tourism that include direct, indirect, and tax contributions are rare in wildlife tourism studies, a comparable calculation for shark-diving tourism in the Maldives using similar methods (though excluding staff service charge), estimated total economic benefits at US$77.1 million in 2016 [52]. This substantial difference highlights the particularly high economic contribution of MRW within the Maldives’ tourism sector. The portion of the US$311 million in revenue generated by MRW tourism that stays in-country and is distributed for environmental and biodiversity conservation remains unknown. However, there is disparity between the amount of revenue generated through nature-based tourism in the Maldives and the portion of the revenue that is used by government agencies for conservation [99]. More than half of the funding that supports in-country conservation comes from unstable international sources (US$16.3 million in 2013) [99].

### Scale and potential negative impacts of tourism

While concentrated in certain regions (i.e., Alif Alif/Dhaal, Baa, Lhaviyani, Kaafu) [52], diving tourism, particularly MRW, has undergone rapid expansion across the Maldives [47,80,103] raising concerns about the sustainability and management of this industry. In 2008, the Maldives had 101 MRW sites (dive sites *n*=91, snorkel sites *n*=10), representing nearly half (48%; 91 of 190 sites) of all known global MRW sites at the time [3]. By 2017, the amount of recognised sites surged to 273 [56]. Our 2021 findings further demonstrate continued growth, with MRW offered by 80% of tourism bases and present in 95% of administrative regions. Tour operators in this study identified a total of 92 sites they considered top five for diving (*n*=65) and snorkelling (*n*=56) with manta rays. Some sites in the regions surrounding the central Malé City received a considerably high portion of the tourism pressure from diving, snorkelling and vessels (i.e., Hanifaru in Baa, Lankan Beyru in Kaafu, Sola Corner in Raa, and Rangali Madivaru in Alif Dhaal). As the top visited site in 2021, the >11,000 annual visitors to Hanifaru generated US$1,514,400 in revenue by tour operators and the Baa Atoll Biosphere Reserve Office. Although Hanifaru does fall within an MPA and is actively monitored and mitigated by regulatory components [38], it does not exclude this site from the potential adverse impacts of wildlife tourism activities on the focal species and their associated habitats [67,79,104].

Cetaceans have shown to react to tourist vessels with active avoidance strategies [105,106]. Such repeated disturbances even lead to displacement from a preferred habitat and reduced fitness at the population level for bottlenose dolphins (Tursiops *sp.*) [105]. In the Maldives, sublethal injuries [68] and tourist behaviour [67] negatively impact manta rays and lead to a possible reduction in reproductive fitness (N. Froman unpublished data). When tourism is unmanaged or is beyond the carrying capacity of an area, the industry is unsustainable and requires further regulation and enforcement. These impacts can become more pronounced during the climate crisis due to a reduced availability of the manta ray’s planktonic prey which is required to sustain reproductive health and overall fitness [60]. These projected reductions highlight the importance of safeguarding known foraging areas and other critical habitats. Despite the expansive nature of MRW activity, the visitation density at these numerous sites remains unquantified. Investigating this crucial metric in the future is essential for robustly assessing and managing potential negative impacts of tourism.

### Management and sanctuaries

There has never been a targeted commercial fishery for manta rays in the Maldives; however, these populations are still vulnerable to bycatch, illegal fishing, and other human-induced stressors [38,68]. The recognition by the Maldives’ government of the economic value of iconic animals such as manta rays, whale sharks, and reef sharks, was used to develop legal frameworks and policies such as the Hanifaru MPA, and the protection of shark and ray species [37,38]. However, successful conservation relies on community support; a lack of buy-in and top-down conservation approaches can lead to non-compliance or policy reversal [107]. Recent attempts to reestablish longline fisheries and legalise shark fishing in the Maldives in 2024 threatened the sanctuary status, but were successfully blocked through community pushback and an online poll [108,109]. Despite clear economic arguments for manta ray protection, competing pressures over resource-use will continue to pose threats. This underscores the need for robust conservation strategies that align the Maldives’ national conservation goals with its international commitments to the United Nations Convention on Biological Diversity (CBD) and the Conservation of Migratory Species of Wild Animals (CMS).

Nationwide there are 42 MPAs in the Maldives covering 0.5% (116.3 km^2^) of the 21,596 km^2^ of Economic Exclusive Zone, but Hanifaru remains the only MPA that is actively monitored and managed [38,56,67]. Awarded MPA status in 2009, regulations limit the number of visitors to 80 snorkellers and five boats at one time, SCUBA diving is prohibited (http://en.epa.gov.mv/regulations), and an entrance fee/videographer permit is required for filming with drones and in-water flashlight videography [110]. Regulations also enforce a code-of-conduct on how to responsibly swim with manta rays and require tour guides to hold a valid licence that is issued after the completion of a short-course [67]. In 2012, the entirety of Baa was formally designated by the president as the Baa Atoll Biosphere Reserve, within which Hanifaru MPA is a core protected zone. The profit generated through the Biosphere Reserve supports the Baa Atoll Conservation Fund, with funds used for community education and research [110]. In 2023, a broader conservation guideline for all protected species including manta rays was developed under the Protected Species Regulation (2021/R-25), providing a code of conduct for snorkelers, divers and boats involved in MRW across the country, regardless of the MPA status [111].

Recognising the value of flagship species as the focus for wildlife tourism is becoming increasingly widespread [2–4,15,42,112]. For example, the combined global value of mobulid meat and dried gill plates in 2013 was 100-fold less than that of the revenue generated through MRW tourism [3,113]. Although criticised as being simplistic, another method to demonstrate the disparity between consumptive and non-consumptive use of this resource is the revenue-generating ability of an individual animal over the duration of their lifetime [3,4,42]. *Mobula alfredi* are known to display high site fidelity [114,115] and remain within the Maldives’ national borders for the duration of their lives. Less is known about *M. birostris*, but studies suggest they are less migratory than previously assumed and often have relatively limited home ranges [116]. When the current population estimate for *M. alfredi* (n=3,500) in the Maldives is considered to be interacting with the tourism industry over their ∼40-year lifespan, the population would generate approximately US$8.6 billion (assuming real discount rate of 5%; see [4]). Unsurprisingly, the estimated lifetime value of an individual *M. alfredi* in the Maldives of US$2.6 million in 2021 has grown substantially in the past decade, as it was previously estimated to generate US$382,000 in its lifetime (2008; accounting for the updated life expectancy estimate of 40 years and the additional tourist expenses) [42]. These valuations can provide a relatable and quantifiable figure that can estimate the economic value of iconic species, which can highlight their value in terms of conservation through sustainable management.

### Intrinsic value of manta rays

Tour operators in the Maldives perceived manta rays to be important to the local community and highlighted their cultural significance. Marine animals such as whales have been recognised for their “natural goodness” (i.e., the good of a creature satisfying the necessities of its life form) which is believed to be a large driver of the “save the whales” movement and the reason commercial whaling was halted [117]. Natural goodness also serves as the primary standard for evaluating our treatment of other life forms, in addition to providing ethical grounds for conservation practices [117]. Flagship species, such as manta rays, can provide motivation for conservation actions, thereby supporting food security and climate stability [118]. As MRW tourism is reliant on the quality of the experience, it in turn promotes a healthy marine environment and engages the public and community in conservation. Furthermore, the conservation of flagship species has ripple effects on the broader ecosystem. Consequently, promoting these species is a strategic approach that yields widespread benefits for ecosystem health, enabling the conservation of intact environments, restoration of degraded ones, conserving biodiversity, and achieving sustainable development goals [18].

Wildlife viewing tourism has the potential to provide a range of education and conservation benefits for visitors [119]. Tourists who learnt about the animals and environment from mediated wildlife watching experiences are more likely to make behaviour/lifestyle changes that benefit the environment [119]. These experiences also contribute to pro-environmental attitudes and improved on-site behaviour changes, with some tourists additionally holding longer-term intentions to engage in conservation actions [119]. An educational presentation or briefing was offered by over half of the MRW tour operators in Maldives (61%), thus reaching at least 17,000 MRW guests in 2021. Many of these tourists travelled internationally, therefore, the knowledge shared and potential behavioural changes by these tourists can be tracked outside of the Maldives, likely to Asia and Europe as tourists from India, Russia, Germany and the United Kingdom were consistently the top visitors to the Maldives [47].

### Caveats

The economic estimates in this study are likely to be conservative. The benefit transfer ratios used to calculate the tourist expenses did not account for domestic transport (i.e., sea plane, speed boat, ferry and private yacht are all key modes of transport within the Maldives), souvenirs and other purchases; dissimilarly to other studies (see [3,8,52]). Some of the economic benefits of this industry are not easily quantified and were not considered in this study. This includes the ripple effect that tourism businesses generate in the local economy, known as economic multipliers, from purchasing goods and services, as well as from employed staff who in turn spend their salaries [52,97,120]. This study also did not account for international travel, despite the majority of visitors travelling from overseas (a large proportion traveling long distances) and therefore incurring international travel expenses [47]. Thus, it is likely that MRW tourism in the Maldives also has an impact on the global economy. Furthermore, our data collection took place in 2021, the year following the COVID-19 global pandemic, when tourist arrival in the Maldives (1.3 million) were 29% below pre-pandemic 2019 levels (1.7 million) [47]. Since then, arrivals have considerably rebounded, reaching 2.05 million in 2024 – a 55% increase from 2021 [121]. This upward trend suggests that a current economic valuation of the MRW industry would likely yield even higher figures, but it also amplifies concerns regarding the industry’s carrying capacity and potential environmental pressures.

### Conclusions

Marine wildlife tourism can be a powerful tool for both economic growth and the conservation of the natural environment. This study provides an updated assessment of MRW in the Maldives and shows a substantial decadal growth in the industry’s scale and the flow-on benefits to the community, government and businesses in the Maldives. To maintain the economic integrity of this industry and to avoid ecological degradation and loss of income, effective strategies to manage MRW activities should be implemented across the nation. However, there is currently a disparity between the amount of economic value generated through nature-based tourism and the amount of government expenditure supporting environmental conservation in the Maldives [99]. It is important that national authorities not only recognise the benefits of MRW tourism but also ensure enough revenue is reinvested into biodiversity conservation, including manta rays and their habitats.

The growth in MRW tourism over the past decade, not only in the Maldives, but likely globally, highlights the need for having updated regional and global assessments of this industry. These kinds of evaluations can help support the case for conservation by demonstrating the economic value of these iconic animals, the ecosystems they rely upon, and the benefits of their protection. They can also promote the non-consumptive use of this resource and aid appropriate development of region-specific management strategies. In the face of multiple human impacts threatening manta ray population health and resulting in global population declines [65,69–74,122], it is crucial to safeguard flagship species that are of intrinsic, social and economic importance. Protecting such species not only conserves biodiversity but also fosters community engagement in conservation efforts, leveraging their cultural and economic value to motivate local stewardship. MRW tourism has the potential to continue developing into a long-term profitable and sustainable industry, resulting in sustainable livelihoods and business revenue for future generations in the years to come.

## Acknowledgments

The authors would like to offer an extended thank you to the tour operators in the Maldives who completed the survey and for those involved in the pilot stages. Many tour operator employees show incredible commitment to conserving the marine environment in the Maldives. Thank you to the *Environmental Protection Agency* staff and the *Maldives Manta Conservation Programme* staff who distributed the survey in-person to tour operators, especially Nashwa Ahmed Manik. We would like to extend our gratitude to Mary O’Malley whose guidance in the initial stages provided direction and resources for this study. The research was carried out under the *Environmental Protection Agency* Protected Species Research Permit (*EPA/2021/PSR-M09*) and in accordance with the University of the Sunshine Coast Human Ethics exemption (*E24002*).

## Supporting information

**S1 Appendix – Tour operator survey questions (attached).**

**S1 Table.** Definitions and key terms used throughout this study.

| Term | Abbreviation | Definition | Source |
| --- | --- | --- | --- |
| Geographical atoll |  | Atolls with distinct reef systems. These are grouped into administrative regions. | Ministry of Fisheries & Agriculture [77] |
| Administrative regions |  | Administrative regions (also known as atolls) are used to organise the country for governance. | Ministry of Fisheries & Agriculture [77] |
| Manta ray watching | MRW | Recreational activities of diving, snorkelling, and on-water observation with the intent of viewing manta rays in the wild, consistent with the criteria used in other studies. | O'Malley et al. [3] |
| Manta ray watching site |  | Manta ray sites are areas where manta ray sightings are known to be frequent seasonally, periodically, or year-round. Sites where manta rays were encountered opportunistically, or out of season, were not included as MRW sites because they do not fit the conservative approach of this study. | O'Malley et al. [3] |
| Manta ray watching guest | MRW guest | Are those who participated in a MRW trip, which may include individuals that participated in multiple trips. |  |
| Trip |  | A trip is taken by guests on a vessel to snorkel or dive with manta rays. |  |
| Tour operator |  | Tour operators are dive and excursion activity centres (i.e., excursion centres offer recreational activities including snorkelling trips), as well as liveaboard vessels. Tour operators function on bases (i.e., resorts, liveaboard and community islands). |  |
| Tourism base | Base | Bases (i.e., resorts, liveaboards and community islands) are the actual venues in where tour operators operate their activity centres from. |  |
| Activity centre |  | In-water activities like diving and snorkelling are ran out of activity centres. |  |
| Excursion centre |  | These businesses offer recreational activities including snorkelling trips. |  |
| Inhabited community island | Community (or comm.) island | These are islands where the general population resides (i.e., towns, villages, fishing and farming communities with permanent human habitation). Some of these islands (especially ones where tourism is present) have dive and excursion activity centres. |  |
| Regular Resort Island | Resort | A < - (star) resort that operates under the concept “one-island-one-resort”, however does not meet the luxury standards and therefore has not been awarded a 5-star rating. |  |
| 5-star Luxury Resort Island | 5-star Resort (5* in figures) | A resort that operates under the concept “one-island-one-resort” and does meet the luxury standards and therefore has been awarded a 5-star rating. |  |
| Liveaboard |  | This is a vessel that has been designed for people to live aboard (multiple-day trips). It is generally used for recreational diving expeditions where divers onboard stay on the vessel for the duration of the cruise and use it as a diving support vessel. The guests onboard pay per person; thus, it is not a private charter. |  |
| Private liveaboard | | Similar to a liveaboard, but the guests pay for the boat, not per person; thus, it is a private charter. A multi-day cruise on a private liveaboards generally cost between US\$12,175 – 84,000. Because of these high prices, it was not possible to decipher the cost per dive and thus, cost per MRW dive. | |
| Guesthouse |  | Guesthouses provide accommodation for tourists (domestic and international) and are generally at a lesser cost than a resort. The vast majority do not function as tour operators and outsource snorkelling and diving activities to other businesses (e.g., dive and excursion centres); thus, were excluded |  |

**S2 Table.** Data collection: details collected and sources, with shorthand for equation in data analysis. Adapted from O’Malley et al. [3].

| Details | Shorthand in equation | Collection date | Ministry of Tourism [47,48] | Maldives Bureau of Statistics [90] | Internet research | Tour operator surveys | Ministry of Tourism [95] | Baa Atoll Biosphere Reserve [89] | Other sources* |
| --- | --- | --- | --- | --- | --- | --- | --- | --- | --- |
| Tour operators in the Maldives |  | 2021-2022 | X |  | X | X |  |  |  |
| Tour operator revenue | *TOR* | 2022 |  |  | X | X |  |  |  |
| Activity price (US\$) | *AP* | 2022-2023 | | | X | X | | | |
| Number of guests per trip | *NG* | 2022 |  |  |  | X |  |  |  |
| Number of trips | *NT* | 2022 |  |  |  | X |  |  |  |
| Number of weeks of MRW season | *NW* | 2022-2023 |  |  | X | X |  |  | X |
| Number of opportunities for MRW | *NO* | 2022 |  |  |  | X |  |  |  |
| Manta ray watching sites |  | 2022 |  |  | X | X |  |  |  |
| Tourist expenses | *TE* | 2024 |  |  | X |  |  |  |  |
| Government TAX (US\$) | *TX* | 2022 | | X | | | | | |
| Staff service charge (US\$) | *SSC* | 2022 | | X | | | | | |
| Staff salary revenue (Maldivian and Foreign) (US\$) | *SSR<sub>M or F</sub>* | 2022 | | | | X | | | X |
| Entry fees to MPA (Resident and Visitor) | *EF<sub>R or V</sub>* | 2022 |  |  |  |  |  | X |  |
| Tour guide licence fee (US\$) | *TGL* | 2022 | | | | | | X | |
| Professional partnerships fee (US\$) | *PP* | 2022 | | | | | | X | |
| Videography permit fee (US\$) | *VP* | 2022 | | | | | | X | |
| Gross Domestic Product (US\$) | GDP | 2022-2023 | X | X | | | X | | |
| Manta ray (*M. alfredi* and *M. birostris*) population size | P | 2024 |  |  |  |  |  |  | X |
| Manta ray (*M. alfredi* and *M. birostris*) lifespan | LS | 2024 |  |  |  |  |  |  | X |

\*Other sources include manta ray researchers, logbook data and published literature.

